# Cocaine and Morphine Converge to Disrupt Chloride Homeostasis in Ventral Tegmental Area GABA Neurons

**DOI:** 10.1101/2025.07.22.666200

**Authors:** Anna C. Pearson, Blake A. Kimmey, Madison B. Taormina, William M. Holden, Alexey Ostroumov

## Abstract

Identifying shared neural mechanisms influenced by diverse classes of drugs of abuse is essential for understanding addiction and for developing broad-spectrum treatments for substance use disorders. Previous studies indicate that many drugs of abuse increase dopamine output from the ventral tegmental area (VTA) by altering the balance of excitatory and inhibitory inputs onto dopamine neurons, thereby promoting maladaptive plasticity within reward circuits. Here, we demonstrate in rats that acute injections of morphine and cocaine, but not saline, disrupt chloride homeostasis in VTA GABA neurons. This disruption is characterized by a depolarized GABA_A_ reversal potential, impaired chloride extrusion, and posttranslational downregulation of the potassium chloride cotransporter KCC2. Although previous studies linked drug-induced posttranslational downregulation of KCC2 in the VTA to glucocorticoid receptor activation, we found that a glucocorticoid receptor antagonist did not prevent cocaine- and morphine-induced disruption of chloride homeostasis. Instead, our data show that dopamine receptor activation is both necessary and sufficient for these alterations. Notably, chloride homeostasis remains impaired 30 days after volitional morphine self-administration, indicating long-lasting plasticity. These findings complement previous work on nicotine and alcohol, suggesting a shared mechanism of inhibitory plasticity in the VTA following drug exposure. Given that chloride dysregulation in VTA GABA neurons influences downstream circuit function and promotes maladaptive behaviors associated with drug use, we propose KCC2 as a promising therapeutic target for substance use disorders.

## 1. Introduction

Identifying common neural adaptations induced by different addictive substances is critical for understanding the mechanisms of addiction and for developing broadly effective treatments for substance use disorders. While individual drugs may act through distinct molecular targets, they often converge on shared neural circuits—particularly those governing reward, motivation, and learning. Understanding how addictive substances alter these circuits can reveal universal mechanisms of maladaptive plasticity, offering a unifying framework for intervention strategies that transcend specific substances. Moreover, studying these shared forms of maladaptive plasticity deepens our understanding of how the brain’s learning and memory systems are co-opted and dysregulated by addiction, ultimately guiding the development of treatments that restore healthy circuit function and improve long-term outcomes.

One major hypothesis posits that all drugs of abuse ultimately converge on a common mechanism: increased dopamine release from neurons originating in the ventral tegmental area (VTA), a midbrain region critical for shaping reward-related behaviors^1,2^. While significant research has focused on dopaminergic neurons in the VTA and the plasticity they undergo following drug exposure, much less is understood about local GABAergic neurons. These inhibitory neurons regulate dopamine neuron activity and thus play a crucial role in shaping dopamine output^3^. Further, VTA GABA neurons send long-range projections to other brain areas, where they modulate circuit activity and influence motivated and addictive behaviors^4,5^.

Growing evidence suggests that in VTA GABA neurons, drugs of abuse alter GABA_A_ receptor–mediated synaptic transmission by disrupting intracellular chloride (Cl^-^) homeostasis, which is maintained by the K^+^/Cl^-^ cotransporter KCC2^6–9^. In the VTA, KCC2 expression is confined to GABAergic neurons, with little or no detectable protein in dopamine neurons^10^. In GABA neurons, disruptions in KCC2 function can impair Cl^−^ extrusion, leading to a depolarizing shift in the GABA_A_ reversal potential (E_GABA_). This shift weakens GABA_A_ receptor-mediated inhibition and can even result in paradoxical GABAergic excitation of VTA GABA neurons. Disruption of KCC2, induced by stress, chronic pain, or exposure to addictive drugs, impairs inhibitory control over dopamine neurons and has been shown to contribute to drug intake and addictive behaviors^6–9,11–13^.

Despite previous evidence that nicotine, alcohol, and morphine downregulate KCC2 expression in the VTA, it remains unclear whether Cl^−^ homeostasis is similarly disrupted by all addictive drugs. Furthermore, the mechanisms underlying KCC2 impairment are not fully understood. The VTA is influenced by multiple neuromodulatory systems that are robustly activated by different experiences and contribute to long-lasting behavioral adaptations. While corticosteroid-induced post-translational modifications can reduce KCC2 function following acute stress^6^, studies of drugs of abuse report neuroimmune-related reductions in total KCC2 protein expression^9^. Understanding the precise regulatory pathways underlying drug-induced KCC2 downregulation is critical for elucidating addiction mechanisms and for developing new treatments. Nevertheless, whether pharmacologically distinct drug classes converge on shared mechanisms to disrupt KCC2 function remains an important open question.

In the current study, we demonstrate that substances representing two distinct classes of drugs, opioids and stimulants, produce a common depolarizing effect on E_GABA_ and impair Cl^-^ extrusion in VTA GABA neurons. In contrast to previous studies of chronic drug exposure^7,9^, we find that acute cocaine and morphine disrupt Cl^-^ homeostasis via post-translation modifications in KCC2, rather than through reductions in total protein expression. These changes are mediated by activation of dopamine D1 and D5 receptors, but not by stress hormones. Additionally, we report long-lasting adaptations in VTA GABA neurons following volitional morphine self-administration, suggesting that this mechanism can contribute to the persistence of maladaptive behaviors associated with substance use disorders. Collectively, these findings promote the possibility of targeting KCC2 to treat an array of substance use disorders.

## 2. Methods

### 2.1 Animals

Male Long–Evans rats (Harlan-Envigo, 300–500 g) were housed in a quiet, temperature- and humidity-controlled facility under a 12/12 h light/dark cycle. Rats were group housed and had ad libitum access to food and water. All rats were handled for at least 5 days prior to the start of behavioral testing. All procedures were conducted in accordance with the Georgetown University Institutional Animal Care and Use Committee (IACUC) guidelines.

### 2.2 Surgery

All surgeries were performed under isoflurane gas anesthesia. For VTA GABA neuron identification, GAD-Cre rats were injected bilaterally with AAV5-pCAG-Flex-EGFP-WPRE (Addgene, wPlasmid #51502) in the VTA at the following coordinates: anterior-posterior (AP) = 5.5, medial-lateral (ML) = ±1.0, dorsal-ventral (DV) = -8.1 (Paxinos, 2007).

For self-administration experiments, rats underwent jugular catheterization. A polyurethane catheter was surgically implanted by threading it subcutaneously over the shoulder blade and inserting it into the jugular vein, where it was secured with sutures. The catheter was connected to a mesh back-mount platform (Instech Laboratories), which was positioned beneath the skin between the shoulder blades and sutured in place. To maintain patency, catheters were flushed daily with 0.2mL of timentin (0.93 mg/mL, Fisher Scientific) dissolved in heparinized saline (1% heparin, Med-Vet International) and sealed with aluminum obturators when not in use.

### 2.3 Acute Intraperitoneal Drug Injection Procedures

Acute injections of morphine sulfate (10 mg/kg, Spectrum Chemical and the NIDA Drug Supply Program) or cocaine (10 mg/kg, Sigma-Aldrich) were given 12-15 hours prior to *ex vivo* electrophysiology experiments. Both morphine and cocaine were dissolved in sterile saline. RU486 (40 mg/kg, Sigma-Aldrich) was dissolved in DMSO and SCH23390 (0.5 mg/kg, Sigma-Aldrich) was dissolved in saline. RU486 and SCH23390 were administered i.p. 10 minutes prior to cocaine or morphine injections.

### 2.4 Intravenous Morphine Self-Administration

Seven days after jugular catheterization, rats were trained to self-administer morphine (0.75 mg/kg/infusion) or saline on a fixed ratio 1 (FR1) schedule for up to 12 hours per day during their active cycle: 18:00 to 06:00. To prevent overdose, the session was terminated after animals reached 75 infusions of morphine. For 14 consecutive days, rats received 5-second infusions of morphine or saline upon pressing the active lever, accompanied by cue light illumination, and followed by a 20-second “time-out” period. An inactive lever was also present; presses on it were tabulated but had no consequences.

### 2.5 Electrophysiology

Horizontal slices (220 μm) containing the VTA were cut using a vibratome (Leica Microsystems) from both juvenile and adult Long-Evans rats (21-28 days or 8-10 weeks old for acute injection experiments, 14-16 weeks old for self-administration experiments). Brains were sliced in ice-cold, oxygenated (95% O_2_, 5% CO_2_) high-sucrose artificial cerebrospinal fluid (ACSF) containing: 205.0 mM sucrose, 2.5 mM KCl, 21.4 mM NaHCO_3_, 1.2 mM NaH_2_PO_4_, 0.5 mM CaCl_2_, 7.5 mM MgCl_2_, and 11.1 mM dextrose. Immediately after slicing, brain sections were transferred into standard ACSF consisting of: 120.0 mM NaCl, 3.3 mM KCl, 25.0 mM NaHCO_3_, 1.2 mM NaH_2_PO_4_, 2.0 mM CaCl_2_, 1.0 mM MgCl_2_, 10.0 mM dextrose, and 20.0 mM sucrose. Slices were continuously oxygenated (95% O_2_, 5% CO_2_), maintained at 32°C for 40 minutes, and then left to equilibrate at room temperature for at least an additional hour.

GABAergic neurons located in the lateral VTA were distinguished based on morphological and electrophysiological features consistent with previously established^6,15–17^. These neurons typically exhibit small somatic dimensions (less than 20 μm in diameter) and display a relatively fast spontaneous firing rate exceeding 7 Hz under zero current conditions in the cell-attached mode. Additionally, they show minimal hyperpolarization-activated current (I_h_), with peak amplitudes below 150 pA in whole-cell recordings. In our previous studies, we showed that cells with these properties were consistently tyrosine hydroxylase (TH) negative^6,8^. A subset of GAD-Cre rats injected with Cre-dependent GFP were also used. GFP-positive cells in the lateral VTA of these animals exhibited similar morphological and electrophysiological properties

To record E_GABA_ from VTA GABA neurons, perforated-patch recordings were used. Gramicidin was first dissolved in methanol (10 mg/ml), then diluted into the pipette solution (135.0 mM KCl, 12.0 mM NaCl, 10.0 mM HEPES, 0.5 mM EGTA, 0.3 mM Tris-GTP, 2.0 mM Mg-ATP, pH 7.2-7.3) to a final concentration of 150 μg/ml. Synaptic stimulation was delivered using a bipolar tungsten electrode (World Precision Instruments) positioned 100–150 μm from the recording site. GABA_A_ receptor-mediated inhibitory post-synaptic currents (IPSC) were isolated using 20 μM DNQX, 50 μM AP5, 1 μM CGP55845. IPSCs were evoked and recorded under voltage clamp at various holding potentials, and E_GABA_ was determined from the current-voltage relationship. To prevent action potential generation at depolarized holding potentials, tetrodotoxin (0.5 μM, HelloBio) was added to extracellular solution. For studies assessing the impact of D1/D5 receptor activation on E_GABA_, slices were incubated in SKF81297 (10 μM, Sigma-Aldrich) for 10 minutes.

To assess activity-dependent synaptic depression, whole-cell voltage-clamp recordings were made during repeated 20 Hz synaptic stimulation. The internal solution used for these experiments included: 123.0 mM K^+^ gluconate, 8.0 mM NaCl, 2.0 mM Mg-ATP, 0.2 mM EGTA, 10.0 mM HEPES, and 0.3 mM Tris-GTP (pH 7.2–7.3). GABA_A_-mediated IPSCs were isolated with 20 μM DNQX, 50 μM AP5, and 1 μM CGP55845.

Electrophysiological recordings were acquired using an Axopatch 200B amplifier (Molecular Devices) and digitized at 20 kHz using pClamp 9.2 (Digidata Interface, Molecular Devices). E_GABA_ recordings were low pass filtered at 10 kHz. Data were analyzed offline using Clampfit 11.2.

### 2.6 Western Blots

The VTA was dissected from horizontal brain slices of adult rats, with slice preparation conducted as outlined in the Electrophysiology section. Membrane fractions were isolated using the Mem-PER Plus Membrane Protein Extraction Kit (Model #89842; Thermo Scientific, Rockford, IL, USA). Protein samples (30 μg) in 2.5% 2-mercaptoethanol were separated on a 4%–15% Precast Protein Gel (#4561083; Bio-Rad) and transferred to a nitrocellulose membrane (Bio-Rad). Primary antibodies included rabbit anti-KCC2 antibody (1:400, #07-432; Millipore, Temecula, CA, USA) and mouse anti-GAPDH antibody (1:400, #MAB374; Millipore). Secondary antibodies were goat anti-rabbit IgG (#T2191; Applied Biosystems, Foster City, CA, USA) or goat anti-mouse IgG/IgM (#T2192; Applied Biosystems), with all antibodies diluted in SignalBoost solution (#407207; EMD Millipore, Billerica, MA, USA). Membranes were developed using Tropix CDP-Star solution (T2218; Applied Biosystems) for 5 minutes and then scanned with the Protein Simple FluorChem R chemiluminescence detector. A densiometric analysis was preformed using AlphaView SA software. Optical densities of KCC2-specific bands were measured and normalized to GAPDH as the loading control.

### 2.7 Statistical Analyses

Statistical analyses were performed using GraphPad Prism (version 10). Two-tailed t-tests or one-way ANOVA test were conducted to compare group means for GABA reversal potential. For Western blot experiments, paired t-tests were used to compare protein expression levels between saline- and drug-treated animals from the same cage. Repeated-measures two-way ANOVA analyses were used to evaluate data from repetitive synaptic stimulation assays and morphine self-administration. Statistical significance was defined as p < 0.05 for all comparisons. All data are presented as mean ± SEM.

## 3. Results

### 3.1 Acute cocaine and morphine impair Cl^-^ homeostasis in VTA GABA neurons

E_GABA_ represents the membrane potential at which evoked inhibitory postsynaptic currents (IPSCs) reverse direction from inward to outward. To determine E_GABA_ in VTA GABA neurons, we conducted gramicidin-perforated patch-clamp recordings, which preserve intracellular anion concentrations, and measured GABA_A_ receptor-mediated inhibitory postsynaptic currents (IPSCs) across different membrane potentials (Figure 1A). E_GABA_ was recorded from adult and juvenile, male and female rats which received a single injection of cocaine (10 mg/kg, i.p.), morphine (10 mg/kg. i.p.) or saline 12-15 hours before slice preparation. VTA GABA neurons from cocaine and morphine-treated rats exhibited a significantly more depolarized E_GABA_ compared to saline-treated rats (Figures 1B and 1C).

**Figure 1:**
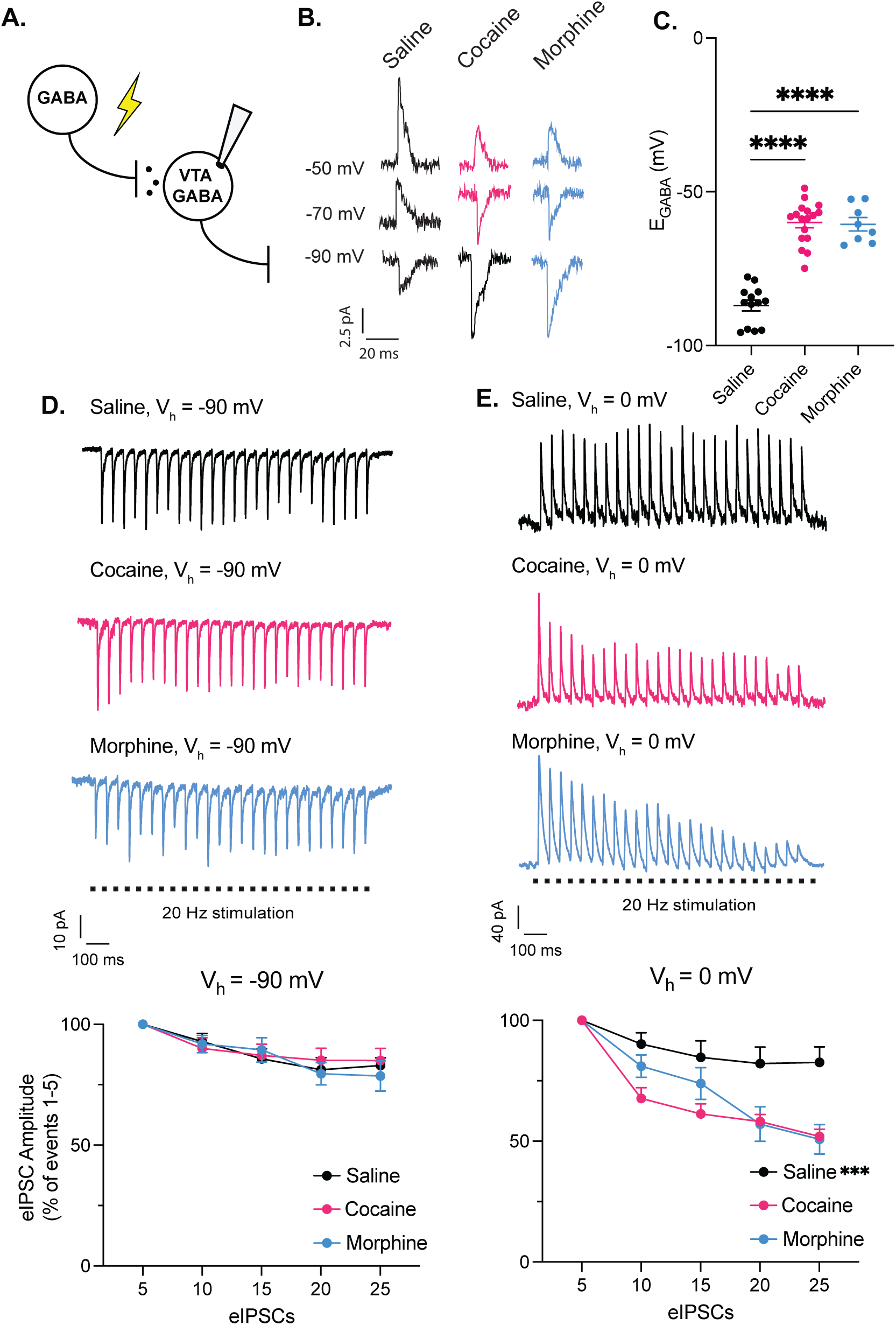
Acute cocaine and morphine impair Cl^-^ homeostasis in VTA GABA neurons. A. To assess cocaine- and morphine-induced alterations in anion homeostasis, GABAergic input onto VTA GABA neurons was recorded using gramicidin-perforated patch-clamp at various holding potentials. IPSCs were elicited through electrical stimulation in the presence of DNQX, AP5, CGP55845, and TTX to isolate GABAergic currents. B. Representative traces of IPSC recordings from saline- (black), cocaine- (pink), and morphine-treated (blue) rats at the given holding potentials. The IPSCs reverse direction at the E_GABA_. The traces were filtered, and stimulus artifacts were removed. C. VTA GABA neurons from cocaine- and morphine-treated rats demonstrated a significantly more positive E_GABA_ value compared to neurons from saline-treated rats (-60.02 ± 1.60 mV after cocaine (pink), and -60.56 ± 2.18 mV after morphine (blue) versus -86.98 ± 1.74 mV after saline (black), *n* = 13 saline, 17 cocaine, 8 morphine cells/group, 7, 5, 7 rats/group, one-way ANOVA, *F_(2, 35)_* = 72.11*, p* < 0.0001). No age nor sex differences were observed in saline and drug-treated rats. D. To assess activity-dependent synaptic depression, whole-cell patch-clamp recordings were performed on VTA GABA neurons during repeated activation of GABA_A_ receptors. Top traces show representative recordings from saline- (black), cocaine- (pink), and morphine-e0xposed (blue) animals, both exhibiting comparable changes in evoked IPSC (eIPSC) amplitude during 20 Hz stimulation at a holding potential of −90 mV. Bottom panel quantifies this effect, showing that the rate of eIPSC amplitude decline did not significantly differ between the cocaine and saline groups (repeated measures 2-way ANOVA, group x time, F_(8, 92)_ = 0.6853, *p* = 0.70, *n* = 9 cells/group, *n* = 4 rats/group). Amplitudes were averaged in sets of five and expressed as a percentage of the mean amplitude of the initial five responses. E. To examine activity-dependent intracellular Cl^−^ accumulation in VTA GABA neurons, a similar protocol was used, but cells were voltage-clamped at 0 mV. Top traces illustrate a modest reduction in eIPSC amplitude in a GABA neuron from a control animal (black) compared to a more pronounced depression in neuron from cocaine- (pink) and morphine-treated (blue) rats. Bottom panel shows that at 0 mV, GABA neurons from cocaine- and morphine-exposed animals exhibited a significantly enhanced synaptic depression relative to saline-treated controls (repeated measures 2-way ANOVA, group x time, F_(8, 92)_ = 4.935, *p* < 0.0001, *n* = 9 cells/group, *n* = 4 rats/group).

Depolarizing shifts in E_GABA_ result from elevated intracellular Cl^−^ levels, often due to impaired Cl^−^ extrusion. During prolonged GABA_A_ receptor activation, reduced Cl^−^ extrusion leads to intracellular Cl^−^ accumulation, ultimately weakening synaptic inhibition by diminishing Cl^−^ gradient. To assess whether acute cocaine or morphine exposure compromises Cl^−^ extrusion in VTA GABA neurons, we applied repetitive GABA_A_ receptor stimulation and measured activity-dependent IPSC depression^6,18,19^. The rate of IPSC amplitude reduction at 0 mV, where Cl^−^ influx dominates, reflects Cl^−^ accumulation and activity-dependent synaptic depression. In contrast, at −90 mV, where Cl^−^ efflux prevails, synaptic depression occurs without accompanying Cl^−^ accumulation.

Following 20 Hz electrical stimulation at −90 mV, cocaine and morphine had no effect on the rate of synaptic depression in VTA GABA neurons (Figure 1D). However, at 0 mV, IPSC amplitude declined significantly faster in cocaine-treated animals compared to saline-treated controls (Figure 1E). The distinct effects of acute cocaine and morphine exposure at −90 mV versus 0 mV indicate increased intracellular Cl^−^ accumulation, suggesting a diminished Cl^−^ extrusion capacity in VTA GABA neurons.

### 3.2 Acute cocaine and morphine decrease KCC2 phosphorylation at Ser940 in VTA

Drug-induced disruption of Cl^-^ homeostasis have previously been linked to reduced expression of the K^+^, Cl^-^ co-transporter KCC2^6–9,13,20^. However, stress-induced depolarizing shifts in E_GABA_ within the VTA have been associated with the decreased phosphorylation at serine 940, rather than changes in total KCC2 protein expression. To investigate whether acute cocaine and morphine administration alter KCC2 expression or phosphorylation within the VTA, we performed Western blot analyses using antibodies against total KCC2 and phosphorylated KCC2 at serine 940 (pS940-KCC2). Immunoblotting revealed prominent bands at 140 and 270 kDa for both total and pS940-KCC2, corresponding to the presence of monomeric and dimeric forms of KCC2 (Figures 2A and 2D). Cocaine exposure did not significantly change total KCC2 levels (Figure 2B). However, the ratio of pS940-KCC2 to total KCC2 was significantly reduced in cocaine-treated animals (Figure 2C). Additionally, we did not find a significant difference in total KCC2 protein expression after morphine exposure (Figure 2E), although the ratio of phosphorylated S940-KCC2 to total KCC2 was significantly reduced following the acute morphine injection (Figure 2F). These findings suggest a post-translational mechanism underlying the observed functional impairment of VTA GABA cells following acute cocaine and morphine exposure.

**Figure 2:**
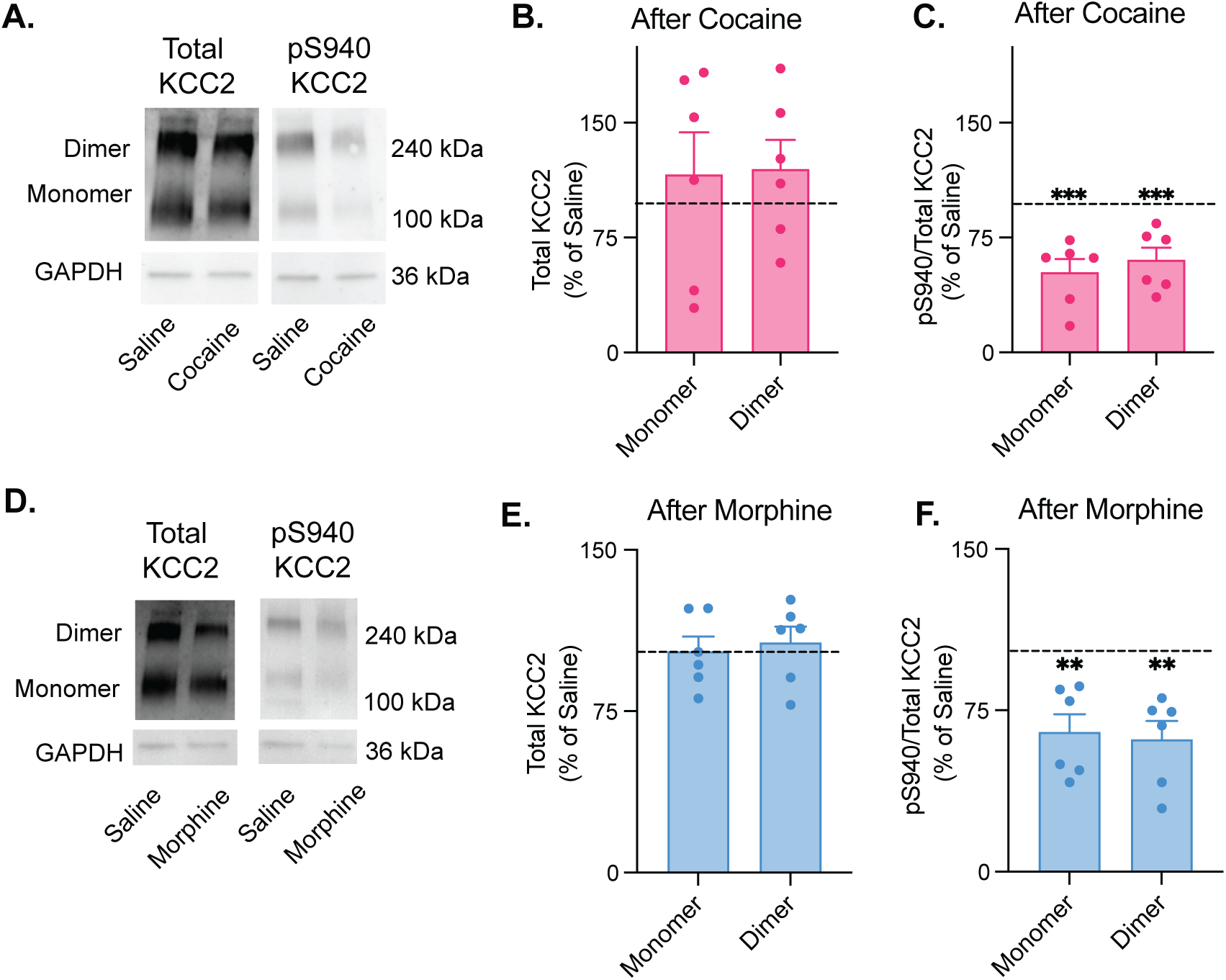
Acute cocaine and morphine decrease KCC2 phosphorylation at Ser940 in VTA. A. Western blot analysis was performed to assess total KCC2 and phosphorylated-S940 KCC2 levels, using GAPDH as a loading control. A representative blot revealed no changes in total KCC2 expression following cocaine. However, cocaine-treated animals exhibited a reduction in pS940 KCC2 relative to total KCC2 compared to saline-treated controls. B. Densitometric analysis did not reveal a significant decrease in total KCC2 expression in cocaine-treated animals compared to saline-treated controls (horizontal dashed line at 100% represents the control mean): 116.0% ± 27.64%, *p* = 0.59 for monomers, 119.6% ± 19.19%, *p* = 0.35 for dimers, *n* = 6 rats/group, normalized to saline controls. C. Densitometric analysis showed a significant decrease in the pS940 KCC2 to total KCC2 ratio in cocaine-treated animals compared to saline-treated controls (horizontal dashed line): 52.28% ± 8.76%, *p* = 0.0028 for monomers, 60.31% ± 8.15% for dimers, *p* = 0.0046, *n* = 6 rats/group, normalized to saline controls. D. Immunoblotting was used to evaluate expression levels of total KCC2 and its phosphorylated form at serine 940 (pS940), with GAPDH serving as the loading control. Representative blots showed that total KCC2 protein levels remained unchanged following morphine exposure. In contrast, morphine-treated animals displayed a decreased ratio of pS940 to total KCC2 compared to saline controls, indicating reduced phosphorylation of KCC2 at this regulatory site. E. Quantification of band intensity revealed no significant difference in total KCC2 protein levels between morphine- and saline-treated groups (horizontal dashed line; 102.7% ± 6.97%, *p* = 0.71 for monomers, 106.8% ± 7.57%, *p* = 0.41 for dimers, *n* = 6 rats/group, normalized to saline controls). F. In contrast, morphine exposure led to a significant reduction in the ratio of phosphorylated S940-KCC2 to total KCC2, as determined by densitometric analysis (horizontal dashed line; 64.81% ± 8.44% for monomers, *p* = 0.0087, 61.47% ± 8.54% for dimers, *p* = 0.0063, *n* = 6 rats/group, normalized to saline controls).

### 3.3 Cocaine and morphine-mediated E_GABA_ depolarization in VTA GABA neurons is not mediated through stress hormones

In the VTA, drug-induced dephosphorylation of KCC2 at serine 940 was previously associated with stress^6–8^, and both acute cocaine and morphine has been previously shown to increase stress hormones^21–24^. Hence, we next investigated whether cocaine or morphine-induced KCC2 downregulation was mediated by stress signaling. Specifically, we assessed whether cocaine- and morphine-induced E_GABA_ depolarization could be blocked by the glucocorticoid receptor antagonist RU486, which has been shown to prevent stress and nicotine-induced impairment of Cl^-^ homeostasis^6,7^.

We performed systemic injections of RU486 10 minutes prior to cocaine or morphine exposure and measured E_GABA_ 12-15 hours after (Figure 3A). RU486 did not block drug-induced depolarizing shift in E_GABA_ in VTA GABA neurons (Figure 3A and 3B), suggesting that the observed effects are not mediated by stress-related pathways.

**Figure 3:**
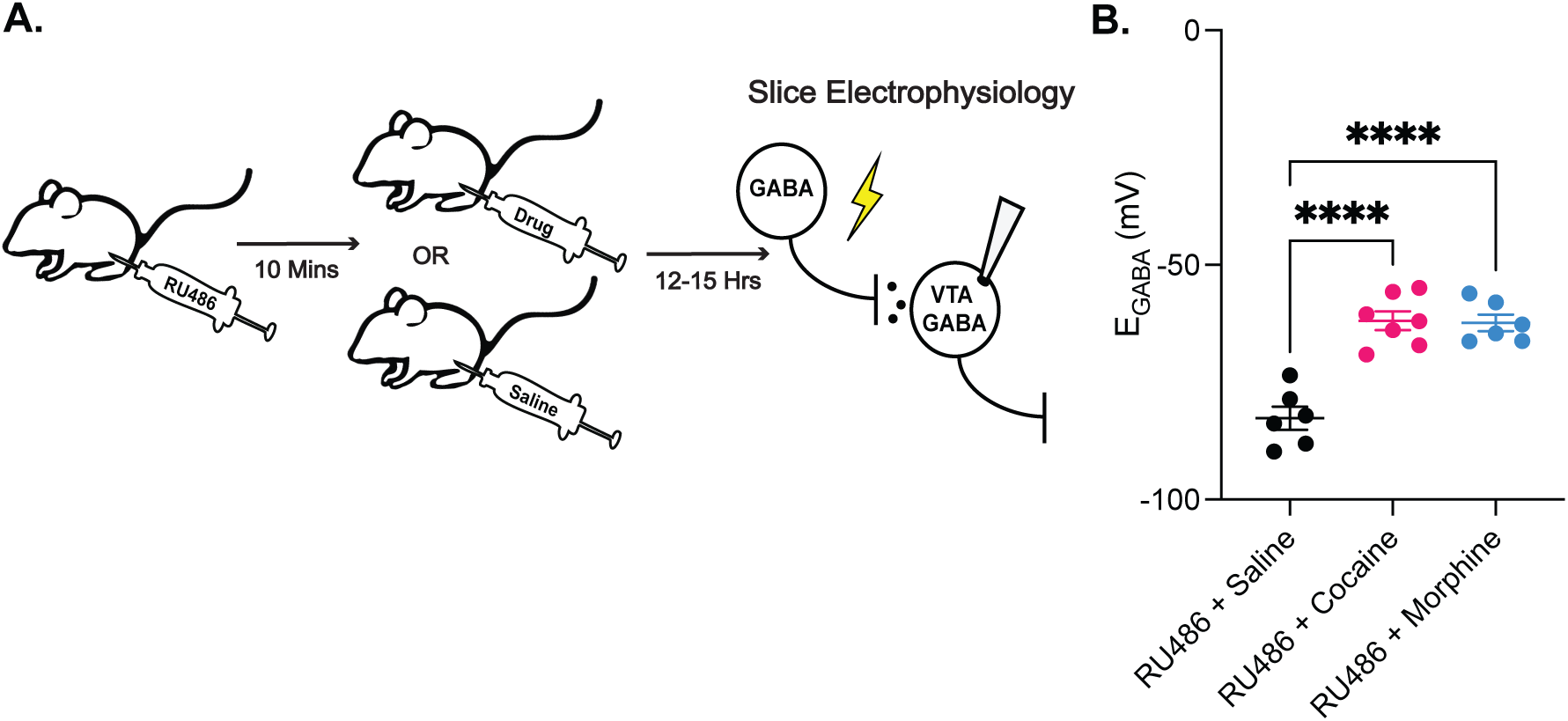
Cocaine and morphine-mediated E_GABA_ depolarization in VTA GABA neurons is not mediated through stress hormones. A. Rats received an i.p. injection of RU486, a glucocorticoid receptor antagonist, 10 minutes before receiving a second i.p. injection of either cocaine, morphine, or saline. 12-15 hours later, rats were sacrificed, and brains were prepared for slice electrophysiology. B. Injections of RU486 did not prevent the depolarization of E_GABA_ in VTA GABA cells in cocaine or morphine treated animals versus saline treated animals: -61.89 ± 2.03 mV for RU486 + cocaine [n = 7 cells, 4 animals; pink data], -62.32 ± 1.77 mV for RU486 + morphine [n = 6 cells, 4 animals; blue data], versus -82.65 ± 2.47 mV for RU486 + saline [n = 6 cells, 4 animals; black data], one-way ANOVA, *F_(2, 16)_*= 30.98, *p* < 0.0001.

### 3.4 Cocaine and morphine-mediated E_GABA_ depolarization in VTA GABA neurons is D1/D5 receptor-dependent

Besides activation of stress system, another shared characteristic of both cocaine and morphine is their ability to elevate dopamine levels within the brain’s reward circuitry^25,26^. Because dopamine release acting through D1 and D5 receptor activation facilitates synaptic plasticity in the VTA^27,28^, we next investigated whether D1/D5 receptor activation plays a role in the depolarizing shift of E_GABA_. First, we systemically injected the selective D1/D5 receptor antagonist SCH23390 10 mins before cocaine or morphine exposure. Then, E_GABA_ was recorded 12-15 hours after drug exposure (Figure 4A). While SCH23390 alone did not alter E_GABA_ in VTA GABA neurons, it completely blocked cocaine- and morphine-induced depolarizing shift in E_GABA_ (Figure 4B).

**Figure 4:**
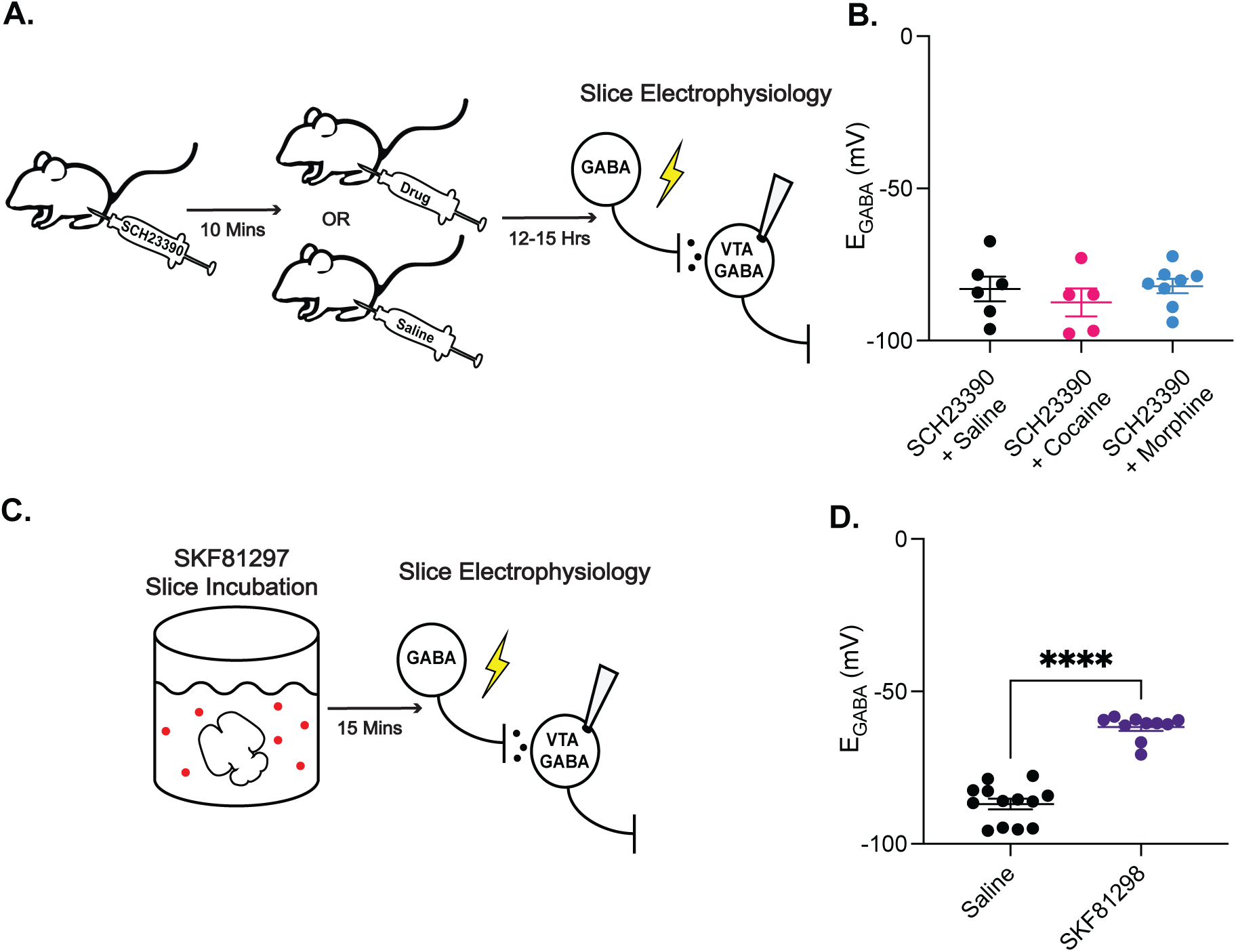
Cocaine and morphine-mediated E_GABA_ depolarization in VTA GABA neurons is D1/D5 receptor-dependent. A. Rats were administered an i.p. injection of the selective D1/D5 receptor antagonist SCH23390 followed 10 minutes later by a second injection of either cocaine, morphine, or saline. After 12–15 hours, animals were euthanized, and brain tissue was collected for *ex vivo* slice electrophysiology. B. Injections of SCH23390 prevented the depolarization of E_GABA_ in VTA GABA cells in cocaine and morphine treated animals versus saline treated animals: -87.46 ± 4.56 mV for SCH23390 + cocaine (n = 5 cells, 2 animals; pink data), -82.10 ± 2.37 mV for SCH23390 + morphine (n = 8 cells, 4 animals; blue data), versus -83.02 ± 4.07 mV for SCH23390 + saline (n = 6 cells, 4 animals; black data), one-way ANOVA, *F_(2, 16)_*= 0.61, *p* = 0.56. C. Brain slices from naïve rats were incubated in the D1/D5 receptor agonist SKF81297 for 15 minutes before cells were recorded. D. Slice incubation SKF81298 resulted in the depolarization of E_GABA_ in VTA GABA neurons in naïve animals; -61.68 ± 1.24 mV for SKF81297-treated cells versus -88.65 ± 3.16 mV (n = 10 cells, 6 animals; purple data) for control cells (n = 13 cells, 7 animals; black data), unpaired t-test, *p* < 0.0001.

Next, we incubated brain slices from drug-naïve animals in the D1/D5 receptor agonist SKF81297 for 15 minutes before recording from VTA GABA neurons (Figure 4C). In contrast to control slices, incubation of brain slices from drug-naïve animals with SKF81297 mimicked the effects of cocaine and morphine, inducing a comparable depolarization of E_GABA_ in VTA GABA neurons (Figure 4D). Overall, these results demonstrate that D1/D5 receptor activation is necessary and sufficient for the cocaine-and morphine-mediated E_GABA_ depolarization in VTA GABA neurons.

3.5 *Morphine self-administration leads to a long-term disruption of Cl^-^ homeostasis*

Compared to passive, non-contingent drug exposure, volitional drug intake can induce differential changes within the mesolimbic system^29–34^. To investigate the potential differences between experimenter-induced and volitional morphine on Cl^-^ homeostasis in the VTA, we used an operant intravenous self-administration paradigm. Rats were trained to self-administer either morphine or saline for 14 consecutive days. Thirty days after the final self-administration session, animals were sacrificed and acute brain slices containing the VTA were prepared for whole-cell electrophysiological recordings to assess changes in GABAergic signaling (Figure 5A).

**Figure 5:**
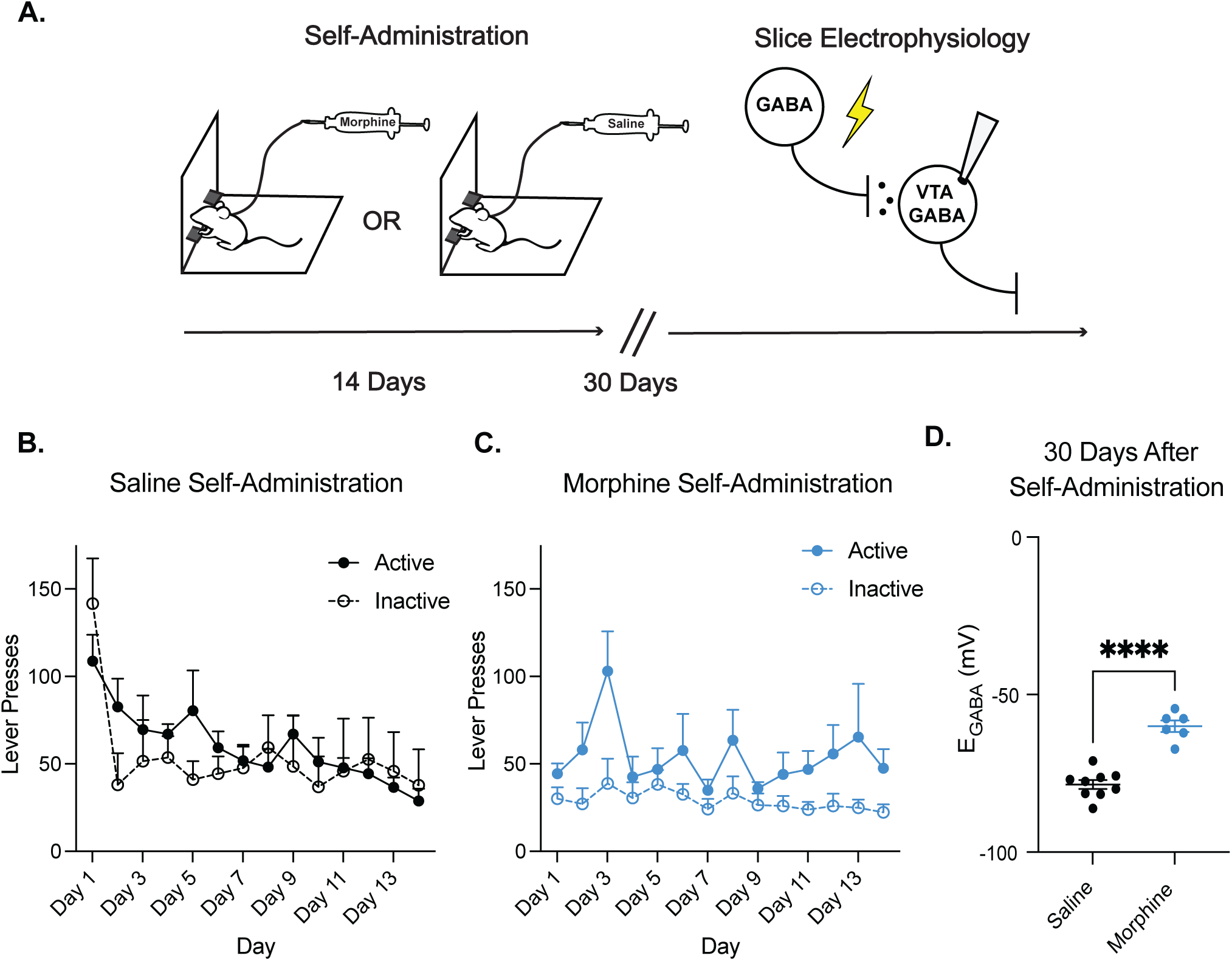
Morphine self-administration leads to a long-term disruption of Cl^-^ homeostasis. A. Rats were trained to self-administer morphine (0.75 mg/kg/infusion) or saline for 12 hours per day, over 14 consecutive days. Thirty days following the final self-administration session, rats were sacrificed for electrophysiology experiments. B. Saline-administering rats (n = 5) did not develop a preference for the active lever over the course of 14 self-administration sessions (two-way ANOVA, effect of group, *F_(1, 8)_* = 0.16, *p* = 0.6999). C. Morphine self-administering rats (n = 9) demonstrated a preference for the active lever (two-way ANOVA, main effect of group, *F_(1, 20)_* = 9.38, *p* = 0.0061). D. Morphine self-administration resulted in a long-lasted depolarization of E_GABA_ in VTA GABA neurons compared to saline self-administration: -60.01 ± 1.83 mV for morphine self-administering animals (n = 6 cells, 2 animals; blue data) versus -78.53 ± 1.44 mV for saline self-administering animals (n = 8 cells, 3 animals; black data), unpaired t-test, *p* < 0.0001.

Rats trained to self-administer saline showed a comparable number of presses on the active and inactive levers (Figure 5B). In contrast, rats trained to self-administer morphine demonstrated a preference for the active lever compared to the inactive lever (Figure 5C). Further, rats self-administering morphine consumed on average 22.08 ± 2.30 mg/kg of the drug per session. Thirty days after final self-administration session, VTA GABA neurons from morphine but not saline animals showed depolarized E_GABA_ (Figure 5D). These findings indicate a persistent disruption of Cl^-^ homeostasis following volitional morphine exposure.

## 4. Discussion

The present study identifies a convergent neural mechanism by which distinct classes of drugs of abuse, specifically cocaine and morphine, impair GABA_A_ receptor transmission and trigger inhibitory plasticity within VTA GABA neurons. Our findings demonstrate that both acute cocaine and morphine exposure lead to a common impairment of Cl^-^ homeostasis in VTA GABAergic neurons, characterized by a depolarized GABA_A_ reversal potential (E_GABA_) and a diminished capacity for Cl^-^ extrusion. Crucially, we show that this disruption is mediated through dopamine D1/D5 receptor activation and involves a decrease in KCC2 phosphorylation at serine 940. Furthermore, we report that volitional morphine self-administration results in a long-lasting impairment of Cl^-^ homeostasis in these VTA GABA neurons. These results underscore the critical role of KCC2 in maintaining inhibitory control in the VTA and suggest its modulation as a novel therapeutic target for substance use disorders

This study is the first to show that cocaine impairs Cl^-^ extrusion in VTA GABA neurons via KCC2 dephosphorylation. However, previous studies have reported that repeated morphine injections lead to impaired GABA_A_ receptor inhibition in VTA GABA neurons^35^ and BDNF-dependent reduction in KCC2 expression^9^. Until the current study, however, the impact of acute exposure to morphine on KCC2 was unknown, as well as the mechanisms behind KCC2 changes. Here, we found that a single injection of either cocaine or morphine was sufficient to impair KCC2 function, as evidenced by a depolarized E_GABA_ and an impaired ability to extrude Cl^−^. While total KCC2 protein expression was not significantly changed, there was a significant reduction in the ratio of phosphorylated KCC2 at serine 940 to total KCC2, implicating a post-translational mechanism in the observed functional impairment. Phosphorylation at serine 940 is known to stabilize KCC2 at the plasma membrane, and its dephosphorylation—mediated by calcium-dependent protein phosphatase 1—promotes KCC2 internalization and functional downregulation^36^. However, it is important to acknowledge that other KCC2 regulatory mechanisms, including alternative phosphorylation sites, glycosylation, and non-coding RNAs, as well as the involvement of other Cl^−^ transporters, may also contribute to the disruption of Cl^−^ homeostasis induced by cocaine and morphine^36, 37^.

Previous studies have shown that acute stress downregulates KCC2, increasing vulnerability to alcohol consumption^6^. Similarly, nicotine exposure has been reported to produce KCC2 downregulation by acting through stress hormones^7^. Our results, however, show that blocking stress hormones, via RU486 administration does not influence cocaine or morphine-induced depolarization of E_GABA_. This finding implies that not all addictive drugs require glucocorticoid release or activation of the hypothalamic–pituitary–adrenal (HPA) axis to disrupt Cl^−^ regulation. Instead, we found that activation of dopamine D1 and D5 receptors was both necessary and sufficient to cause E_GABA_ depolarization following cocaine and morphine exposure implicating a previously unrecognized dopaminergic mechanism in KCC2 downregulation. While our pharmacological manipulations implicate dopamine and rule out glucocorticoids, other neuromodulatory systems activated by cocaine and morphine—such as serotonergic, noradrenergic, or opioid signaling—may also contribute to KCC2 modulation and remain to be investigated.

Because prior studies examining KCC2 in the context of opiates have relied on passive drug administration models^9,20^, we sought to determine whether volitional drug intake—known to engage distinct neural adaptations^30,34^—would differentially alter E_GABA_. Unlike passive drug delivery, self-administration more closely mimics the patterns and neurobiological consequences of human opioid use, including the development of drug-taking habits, altered reward processing, and persistent adaptations in mesolimbic circuits. Our findings indicate that 2-week morphine self-administration leads to a lasting disruption of Cl^−^ homeostasis in VTA GABA neurons, potentially contributing to the neural adaptations underlying persistent addictive behaviors. This sustained impairment parallels the long-lasting alterations in VTA GABAergic signaling observed after adolescent nicotine exposure, but contrasts with the transient effects of acute or two-week experimenter-administered nicotine in adulthood, which typically subside within days to a few weeks^7,38^. The molecular basis for the long-term Cl^−^ dysregulation observed after morphine self-administration remains undefined. Although acute and chronic effects likely involve overlapping mechanisms, the persistence of these adaptations suggests involvement of transcriptional or epigenetic reprogramming, processes that were not examined in the current study.

By identifying a convergent mechanism of inhibitory dysfunction, these findings highlight KCC2 as a promising molecular entry point for restoring normal VTA circuit function across multiple drug classes. Notably, reduced KCC2 function in VTA GABA neurons may shape behavior by altering the activity of distinct dopamine and GABA output pathways. To date, this downregulation has been shown to reduce drug-evoked dopamine release within the mesolimbic VTA-NAc pathway, a circuit critical for reward and addictive behaviors^6–9,13,38^. Given the functional diversity of VTA DA projections^39–43^, future studies should explore whether KCC2 dysregulation differentially affects distinct mesolimbic subcircuits, which have dissociable roles in reward-related behaviors^44,45^. In addition to mesolimbic circuits, KCC2 downregulation may impact VTA dopamine projections to cortical and amygdala regions, which are also implicated in addiction^4^. Furthermore, KCC2 can be downregulated in VTA GABA projection neurons, which directly target forebrain and brainstem regions involved in reward and aversion^46–49^. Linking these cellular adaptations to changes in dopamine and GABA signaling, motivation, or drug-seeking behavior will be a crucial next step in establishing the functional relevance of KCC2 downregulation. Thus, future studies aimed at defining behavioral consequences of KCC2 dysregulation may provide important insights for developing interventions that reverse the maladaptive adaptations underlying addictive behaviors.

